# Autonomic responses to proprioceptive and visual errors during single-trial reach adaptation

**DOI:** 10.1101/2025.03.31.646172

**Authors:** Motoko Nogami, Nobuhiro Hagura, Atsushi Yokoi

## Abstract

The autonomic nervous system (ANS) plays a crucial role in coordinating brain and bodily responses to cope with unexpected events. Extensive research in cognitive control has demonstrated that error-induced responses in peripheral autonomic measures, such as pupil diameter and heart rate, correlate with the quality of the subsequent behavioral adjustment. Here, we perform ANS measurements during a motor learning task to investigate error-induced ANS responses in implicit motor learning. In a series of experiments, we measured multimodal autonomic signals including pupil diameter, skin conductance, and heart rate while participants made a reaching movement under occasional proprioceptive or visual perturbations of different magnitudes. We characterized the phasic autonomic responses to the induced errors and evaluated their influence on motor learning, which was quantified by comparing the trials immediately before and after the error event (single-trial reach adaptation). Consistent with previous research, the results revealed the sequence of ANS responses induced by unexpected motor errors; the pupil dilation, increased skin-conductance, and heart rate deceleration. These responses also demonstrated a clear dose-dependent increase in responses to errors in both visual and proprioceptive modalities. Furthermore, a latent factor analysis using motor learning and multimodal autonomic response data allowed us to detect a statistical relationship between the latent autonomic states and the motor learning rate, suggesting the suppression of implicit motor adaptation by error-induced sympathetic activity. These results provide novel insights into how ANS-tagged internal states affect implicit motor learning.

**Significance statement:** The autonomic nervous system (ANS) plays a crucial role in coordinating brain and bodily responses when coping with unexpected situations. Although extensively studied in the field of cognitive control, little is known about whether and how ANS is involved in implicit motor learning. Here, by monitoring multimodal ANS signals (pupil, skin conductance, and heart rate) during a motor adaptation task, we show that ANS measures show a size-dependent phasic response to proprioceptively/visually applied motor errors. We further show that the ANS, especially the sympathetic error response, mediates the suppression of error sensitivity in larger errors. These results provide novel insights into how ANS-tagged internal states, potentially relevant for statistical inference, affect implicit motor learning.

**Supplementary Materials:** One Supplementary Material file is available.

## Introduction

The autonomic nervous system (ANS) plays a crucial role in coordinating brain and bodily responses to cope with unexpected, challenging situations such as seeing a bear while hiking. In such situations, a close interplay among different brain regions is critical to rapidly assess one’s external/internal states and make a swift decision to act (or not) (Roelofs and Dayan, 2022). For example, the detection of an unexpected event in medial prefrontal cortical areas, such as the anterior cingulate cortex (ACC) (Critchley et al., 2003, 2005), triggers the activation of subcortical areas, such as the locus coeruleus (LC), which then interrupt the ongoing mental processes through noradrenaline (NA) and reallocate attentional resources for necessary cognitive processes (Bouret and Sara, 2005; Sara, 2009; Sara and Bouret, 2012). These brain responses are paralleled by bodily responses mediated by the ANS, such as increased heart rate, sweating, and pupil dilation, as well as hormonal release from the adrenal medulla, to prepare for the upcoming mental/physical challenge [i.e., “ fight or flight” (Canon, 1915)].

Although not as life-threatening as a bear, similar processes are observed when people face an error in tasks that involve cognitive control and/or decision-making. Error-related ANS responses such as pupil dilation and heart rate deceleration in cognitive tasks have been shown to predict the quality of the behavioral adjustment in following trials (Hajcak et al., 2003; Preuschoff et al., 2011; Wessel et al., 2011; Nassar et al., 2012; Murphy et al., 2016). These ANS responses are therefore regarded as a peripheral readout of the cognitive control process triggered by an error. Interestingly, while ANS responses have been extensively investigated as indices for cognitive learning, the similar contribution of the ANS to implicit motor learning has been less studied. Thus, we recently set out to assess pupil diameter changes during short-term motor adaptation to a novel force field in humans. Although we did not directly assess the relationship between the pupil responses and motor learning quality, the results suggested that the phasic pupil dilation in response to movement errors reflects sensory surprise and the tonic pupil diameter subjective uncertainty regarding the task environment (Yokoi and Weiler, 2022). However, although pupil diameter is regarded as an established measure of the central LC-NA system (Joshi and Gold, 2020), pupil diameter alone does not define a comprehensive ANS state, as it is controlled by the sympathetic and parasympathetic balance (Mathôt, 2018).

In the current study, to obtain a more comprehensive picture of error-induced ANS responses and more directly examine their association with implicit motor learning, we assessed multimodal autonomic signals during a single-trial learning paradigm (Marko et al., 2012; Kasuga et al., 2013; Hayashi et al., 2020; Makino et al., 2023). We measured pupil diameter, skin conductance, and heart rate to assess their responses to differing sizes of unexpected motor errors applied proprioceptively or visually. We also examined the relationship between ANS responses and the learning rate (i.e., sensitivity to errors), quantified by comparing the trials immediately before and after the error event. The results demonstrated a dose-dependent increase in all phasic ANS responses investigated in response to errors in both modalities. Using a latent factor analysis combining motor learning and the multimodal autonomic response data, we confirmed a statistical relationship between the latent autonomic states and the motor learning rate. Our results suggest involvement of the error-driven autonomic state (or LC-NA activity) change in the adjustment of the motor learning rate. Such ANS activity may reflect the involvement of explicit cognitive processes for large detectable errors. Moreover, together with previous studies showing that the phasic ANS response reflects surprise- or uncertainty-like states (Preuschoff et al., 2011; Nassar et al., 2012; de Berker et al., 2016; Zénon, 2019), ANS signals could allow to assess latent computational variables mediated by statistical inference, such as contextual inference (Heald et al., 2023).

## Materials and Methods

### Participants

Eighty-five human volunteers (58 men and 27 women, 19–31 years old) participated in the study. Forty-one individuals participated in Experiment 1 and 44 in Experiment 2. Sample size per experiment was based on a previous study collecting multiple autonomic indices (de Berker et al., 2016). All participants were right-handed, verified with the Flanders Handedness Questionnaire (Nicholls et al., 2013; Okubo et al., 2014), and claimed no history of neurological disorders. The experimental protocols were approved by the ethical committees of the National Institute for Information and Communications Technology. Participants provided written informed consent before participation in the study.

### Apparatus

Participants sat in front of an LCD monitor and grasped the handle of a robotic manipulandum (Phantom Premium HF 1.5, 3D Systems, Inc.) with their right hand. They also rested their chin on a chin rest positioned in front of the monitor. An eye tracker (EyeLink 1000, SR Research Inc., Ontario, Canada) was placed under the monitor to measure the participants’ pupil diameter (Fig. 1A). The chin rest and monitor were 44 cm apart. The coordinates for visual stimuli (start and target positions and a cursor representing the hand position) were located in the x-y plane, while the handle motion was constrained to the x-z plane (Fig. 1A). The colors of visual objects were adjusted to approximately equal luminance to avoid changes in pupil diameter induced by task stimuli (Yokoi and Weiler, 2022). Participants were unable to directly see their hand.

**Figure 1.**
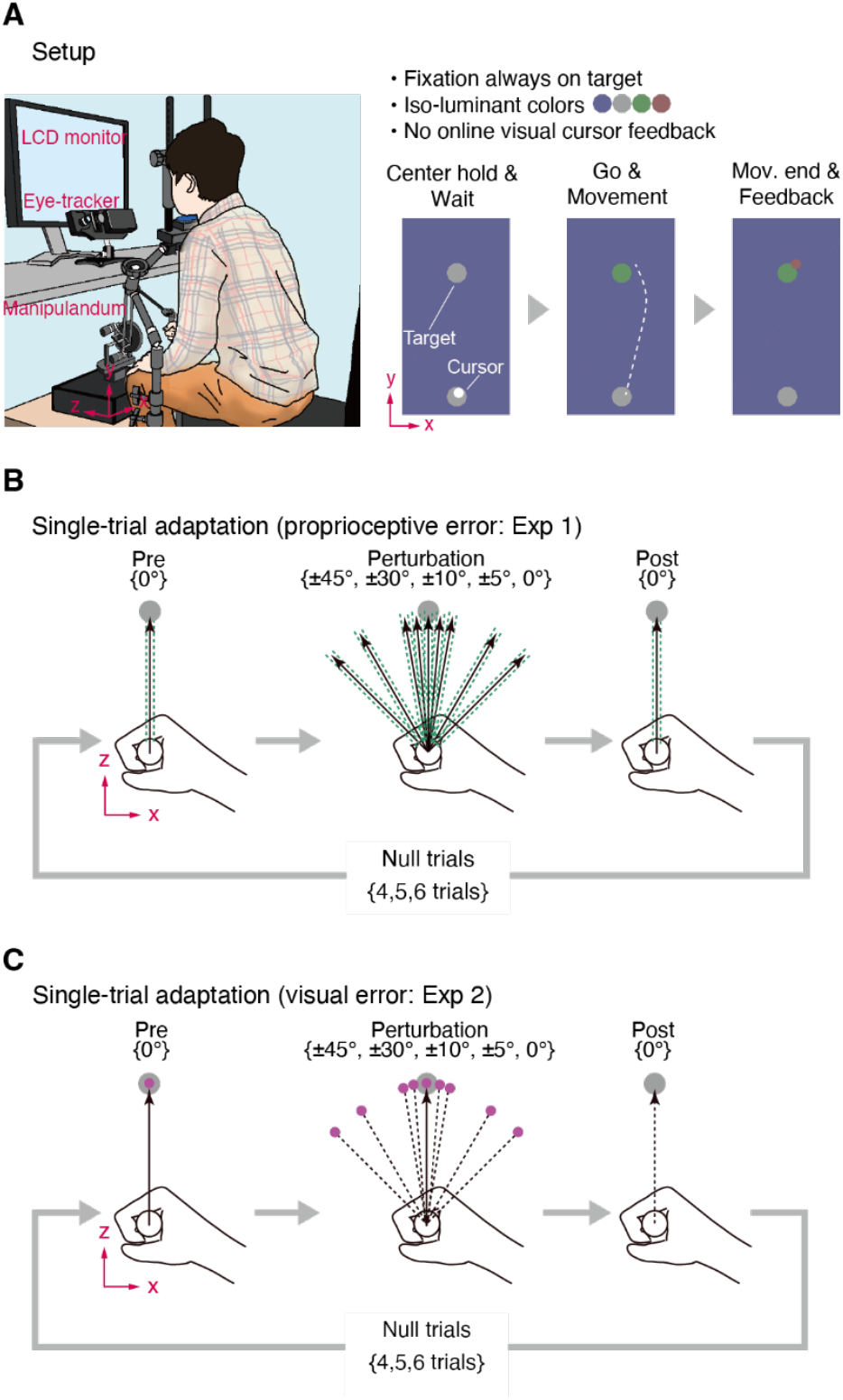
Simultaneous acquisition of multimodal autonomic signals (pupil, skin conductance, and heart rate) during the single-trial motor adaptation paradigm. **(A)** Experimental setup for experiments 1 and 2. We measured the pupil diameter, skin conductance, and electrocardiogram of the participants while they performed the reaching movement holding the handle of the robotic manipulandum (left). Participants initially practiced making straight reaching movements to hit a cursor into a target (right). In the main session, however, the online cursor feedback was occluded, and only terminal feedback was provided. **(B, C)** Single-trial adaptation to various error sizes assessed by comparing the movements in trials immediately before (“ Pre”) and after (“ Post”) the perturbation trial. In Exp. 1, we assessed the adaptation to the proprioceptive error by constraining the reach trajectory in nine different directions with a simulated “ force channel” (B). In pre- and post-perturbation trials, we measured the participants’ force in the direction perpendicular to the movement with the force channel. To minimize the mismatch between visual and proprioceptive information, no endpoint feedback was provided in perturbation trials. In Exp. 2, we assessed the adaptation to visual error by applying visuomotor rotation of the cursor in nine possible angles. To this end, continuous visual cursor feedback was provided but no feedback was provided in post-perturbation trials.

Electrocardiographic signals (ECG) and skin-conductance level (SCL) were measured using the BIOPAC system (BIOPAC MP160, BIOPAC SYSTEMS Inc.). ECG electrodes were placed at three points on the trunk (below the right and left clavicle and on the left rib). In addition, SCL electrodes were attached to the middle and index fingers of the left hand. As an additional index, photoplethysmography (PPG) was collected for 19 participants in Exp 1 and 29 participants in Exp 2. A pulse-oximeter sensor was attached to the left ring finger of each participant. Nonetheless, PPG data were not analyzed due to the insufficient sample size compared to other autonomic signals.

### Procedure

Participants were instructed to move a cursor (a white 5 mm diameter dot) as straight as possible from the starting position (a gray 16 mm diameter dot) to the center of the target (a gray 16 mm diameter dot). After moving the cursor, they were asked to maintain the cursor on target until the end of the trial (Fig. 1A). Home and target positions were 10 cm apart. We instructed the participants to always aim straight at the target. Participants were required to minimize face movements to keep their gaze fixed on the target. They were also asked to keep their left hand still and suppress blinking during the movement to reduce noise in skin-conductance response (SCR) and pupil measurements. Every single-trial started after a stable fixation to the target and handle holding at the starting position for 1.0 s. This was followed by a random delay of 1500–1900 ms before a GO cue, the change in target color to green. In the experiment, the movement onset and offset were defined as the time points when the speed of the handle was >3.5 cm/s or <0.1 cm/s, respectively. Movements requiring >1.5 s to complete were warned by a visual message “ Too Slow!.” After movement offset, endpoint feedback was provided for 3.0 s. This feedback duration was chosen to reliably measure the SCR to movement errors, which typically peaks around 1.5–3.0 s from an event (Dawson et al., 2007). After the feedback period, the manipulandum handle automatically returned to the start position and the trial ended.

To accurately measure ANS error responses and the error-dependent change in reaching behavior (i.e., learning), we employed a single-trial learning paradigm with varying sizes of movement error (Marko et al., 2012; Kasuga et al., 2013; Hayashi et al., 2020). After every 4–6 trials of normal reaching movements without perturbations, a set of three trials was introduced to assess the single-trial motor adaptation. This “ triplet” consisted of two probe trials with one perturbation trial in between. The learning induced by the perturbation was quantified by comparing the motor outputs before and after the perturbation trial. The perturbation size was randomly chosen from the set *p* = {±45°, ±30°, ±10°, ±5°, 0°}.

Before the main session, participants conducted two practice blocks with continuous cursor feedback (30 trials each) in the absence of any perturbations. The main experiment consisted of seven blocks, each containing 72 trials. After each block, there was a 1-min break and after the third block, a 5-min break. After the main experiment, pupil calibration was performed by changing the background color of the monitor from light blue to either white (high luminance) or black (low luminance). This triggered the pupil light reflex, allowing the measurement of each participant’s minimum and maximum pupil size in the current experimental environment. The obtained range of pupil sizes was then used to normalize the pupil data within each participant, ensuring consistent data interpretation across participants (Yokoi & Weiler, 2022). Following this calibration session, participants answered a questionnaire asking whether they noticed any perturbation, whether they could predict the perturbations, and whether they intentionally aimed at nontarget directions.

### Experiment 1 (proprioceptive error)

Forty-one participants were recruited for Exp. 1. Two participants were excluded from the analysis due to technical failure to measure either pupil or SCL/ECG data. As a result, the data of 39 participants were used for further analysis (age [mean ± SD]: 22.5 ± 2.65; 27 male, 12 females; handedness score [mean ± SD]: 9.74 ± 0.60). In Exp. 1, we assessed pupil/autonomic responses to proprioceptive errors during reaching, which were induced by constraining the hand trajectory to a specific direction (±45°, ±30°, ±10°, ±5°, or 0°) by using a force channel method (Scheidt et al., 2000; Hayashi et al., 2020), which mechanically constrains the handle trajectory (Fig. 1B). Similarly, the motor output before and after the perturbation trial were measured by the force channel directed straight to the target (Fig. 1B; 0°). The channel stiffness and viscosity were 23 N/cm and 0.2 N/(cms^-1^), respectively. In Exp. 1, only endpoint feedback was provided (no online cursor feedback). Similarly, endpoint feedback was absent from perturbation trials to minimize the visual-proprioceptive discrepancy (i.e., visual error).

### Experiment 2 (visual error)

Forty-four participants were recruited for Exp. 2. Of those, five participants were excluded from the analysis due to technical failure to measure either pupil or SCL/ECG data. As a result, the data of 39 participants were used for further analysis (age: 22.1 ± 2.16; 28 male, 11 females; handedness: 9.69 ± 0.95). In Exp. 2, we assessed pupil/autonomic responses to visuomotor errors. For consistency with Exp 1, where the error information was available during movement, we provided both online and endpoint cursor feedback (a magenta dot of 7.5 mm diameter) here. The force channel was not used in the entire experiment. Visuomotor errors were applied by rotating the cursor trajectory in a specific direction (±45°, ±30°, ±10°, ±5°, or 0°; Fig. 1C). Similarly, for consistency with Exp 1, endpoint feedback was not provided in perturbation trials. While neither online nor endpoint feedback was provided during post-perturbation trials, it was available during pre-perturbation trials to ensure participants remained unprepared for perturbations.

### Data analysis

All analyses were conducted using custom-written codes in MATLAB R2015b (Mathworks, Natick, MA, USA). For all experiments, data from the manipulandum (x-z positions, x-z velocities, and x-z command forces for the manipulandum handle) and the eye tracker (x-y gaze positions and pupil diameter) were sampled at 200 Hz. Other autonomic data (ECG, SCL, and PPG) were sampled at 1000 Hz.

#### Kinematic data

The kinematic data for arm reaching were first smoothed with a Gaussian kernel of 35 ms full width half maximum (FWHM). Movement onset was re-defined as the point when the hand movement velocity first exceeded a threshold (10% of the peak velocity) in each trial. Similarly, movement offset was defined as the point when the hand movement velocity first fell below the threshold after movement onset.

To assess the single-trial error-dependent change in the motor output and the sensitivity to errors (i.e., learning rate), we employed the following equations:

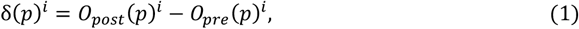

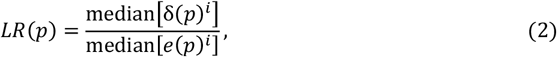

where the *δ*(*p*)^*i*^ represents the change in motor output (i.e., across-trial correction) at the *ith* experimental block (*i* = 1, 2, …, 7) with the perturbation condition *p* (*p* = {±45°, ±30°, ±10°, ±5°, 0°}). Similarly, *O*_*post*_(*p*)^*i*^ and *O*_*pre*_(*p*)^*i*^ represent the motor output for the post- and pre-perturbation trials at the *ith* experimental block with the perturbation condition *p*, respectively. *LR*(*p*) represents the single-trial learning rate (sensitivity to errors) and *e*(*p*)^*i*^ the motor error for the perturbation condition *p* at the *ith* block.

For Exp 1, similar to Hayashi et al. (Hayashi et al., 2020), we employed the force impulse for calculating the *O*_*pre*_, and *O*_*post*_ in Eq. 1. Impulse values were defined as the time integral of the lateral force applied to the channel wall from movement onset to the peak velocity. Similarly, we calculated the force impulse at the perturbation trial as an index for the proprioceptive error (*e* in Eq. 2). The single-trial learning rate (error sensitivity) was calculated as the median of across-trial correction for all blocks divided by the median of proprioceptive errors in all blocks for each perturbation size (Eq. 2). For Exp. 2, we used the hand movement angle for *O*_*pre*_ and *O*_*post*_ in Eq. 1 and defined as the angle relative to the target direction calculated at the time of peak velocity, *T*_*peak*_;

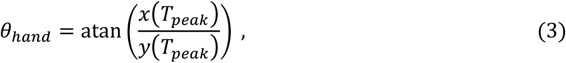

where *x*(•) and *y*(•) represent the *x*- and *y*-positions of the handle relative to the starting position. For the visuomotor error (*e* in Eq. 2), we used the angular deviation of the cursor relative to the target direction (i.e., hand angle + rotation angle). Similar to Exp 1, the rotation angle was randomly chosen from *p* = {±45°, ±30°, ±10°, ±5°, 0°}. The single-trial learning rate (error sensitivity) was similarly calculated as the median of the across-trial correction for all blocks divided by that of the visuomotor errors in all blocks for each perturbation size (Eq. 2).

### Eyetracker data

For all experiments, x-y gaze position and pupil data were preprocessed as follows: first, we discarded the data around each blink event, from 100 ms before and up to 150 ms after the blink. We then interpolated the discarded data using a piecewise cubic Hermite interpolating polynomial (Matlab interp1.m function with “ pchip” option). Next, the gaze position data were smoothed by a second-order Savitzky-Golay filter with a frame length of 55 ms (sgolayfilt.m function). The eye velocities were then derived from the filter results. We employed 30°/s as velocity threshold for saccade detection. For the pupil diameter data, to remove high-frequency noise, we smoothed the data with a Gaussian kernel with 235 ms FWHM. Furthermore, we individually normalized the pupil diameter data relative to the minima and maxima of the pupil diameter data measured during light reflex trials. The normalized pupil diameter data were used for further analyses.

We defined the tonic component of pupil changes as the pupil diameter measured before go-cue presentation. We also defined the phasic pupil response to errors as the pupil dilation velocity after movement onset. The pupil dilation velocity was derived through a numerical derivative (diff.m function). The trial-by-trial summaries of these pupil-related variables were defined as follows: the tonic baseline pupil diameter was defined as the average pupil diameter during the waiting period before the onset of the GO-cue. The trial-by-trial phasic pupil responses to errors were defined as the average pupil dilation velocity within the temporal regions of interest (ROIs) where the significant cluster of perturbation size was detected (Tables 1 and 3, Small vs. Large). This was assessed by comparing the average time series of the pupil dilation velocity at perturbation trials for small (|*p*| ≤ 10°) against large (|*p*| ≥ 30°) errors using the cluster-based permutation test (Maris and Oostenveld, 2007).

**Table 1.** Temporal ROIs for the significant effect of proprioceptive perturbation (Exp 1) tested over the phasic autonomic response time series aligned to movement onset (Fig. 2, left column).

| Variables | Comparison | Start | Center | End | Extent | p-Values |
| --- | --- | --- | --- | --- | --- | --- |
| Pupil velocity | Null vs. Large | 170 | 480 | 790 | 620 | $1.00 \times 10^{-4}$ |
| | | 1510 | 1900 | 2295 | 785 | $1.00 \times 10^{-4}$ |
|  |  | 2940 | 3235 | 3535 | 595 | 0.002 |
|  |  | 1060 | 1225 | 1395 | 335 | 0.002 |
| | Null vs. Small | 2575 | 3050 | 3525 | 950 | $1.00 \times 10^{-4}$ |
|  |  | 455 | 635 | 820 | 365 | 0.003 |
|  |  | 1025 | 1190 | 1360 | 335 | 0.004 |
|  |  | 1545 | 1695 | 1845 | 300 | 0.022 |
| | Small vs. Large | 170 | 470 | 775 | 605 | $1.00 \times 10^{-4}$ |
| | | 1545 | 1900 | 2255 | 710 | $1.00 \times 10^{-4}$ |
|  |  | 1090 | 1225 | 1360 | 270 | 0.014 |
| SCR | Null vs. Large | 1465 | 2455 | 3445 | 1980 | $1.00 \times 10^{-4}$ |
| | Small vs. Large | 1595 | 2545 | 3495 | 1900 | $1.00 \times 10^{-4}$ |
| RRI | Null vs. Large | 1800 | 2705 | 3610 | 1810 | $1.00 \times 10^{-4}$ |
| | Null vs. Small | 2110 | 2775 | 3440 | 1330 | $1.00 \times 10^{-4}$ |

We used a cluster-based permutation test (Maris and Oostenveld, 2007). The overall alpha-level was set to $p < 0.05$ , the cluster-forming threshold to $p < 0.01$ , the test directionality was two-sided, and the permutation number was $10^4$ . Units are in ms. Data within each comparison type are ordered according to the p-values.

### BIOPAC data

SCL data were first filtered with the Second-order Savitzky-Golay filter using a window size of 300 ms. We then defined the SCR as the numerical derivative of SCL data estimated using the Savitzky-Golay filter. Then, both SCL and SCR data were downsampled to 200 Hz for further analyses.

ECG data were filtered using the third-order Savitzky-Golay filter with a window size of 30 ms, and the time derivative of the filtered ECG data were estimated through the filtered result. These data were then used to detect the timing of R-wave peaks. We then estimated the instantaneous R-R intervals using the following equations:

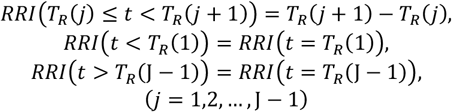

where *T*_*R*_(*j*) represents the time for the *j*-th R-wave peak and J the total number of the detected R-wave peaks at a given trial. Then, the estimated instantaneous R-R interval (RRI) data were smoothed using a Gaussian filter with a 235 ms FWHM and *z*-transformed within each individual.

Similar to pupil data, the trial-by-trial summary for SCL, SCR, and RRI data were defined as follows. The tonic SCL was defined as the average of the SCLL during the pre-movement period. The phasic SCR responses to perturbation were defined as the average values for the temporal ROIs detected by the cluster-based permutation test of the timeseries data for the small against the large perturbation for onset-aligned data (Tables 1 and 3, Small vs. Large). We opted to choose offset-aligned data for phasic RRI responses (Tables 2 and 4, Small vs. Large).

**Table 2.**
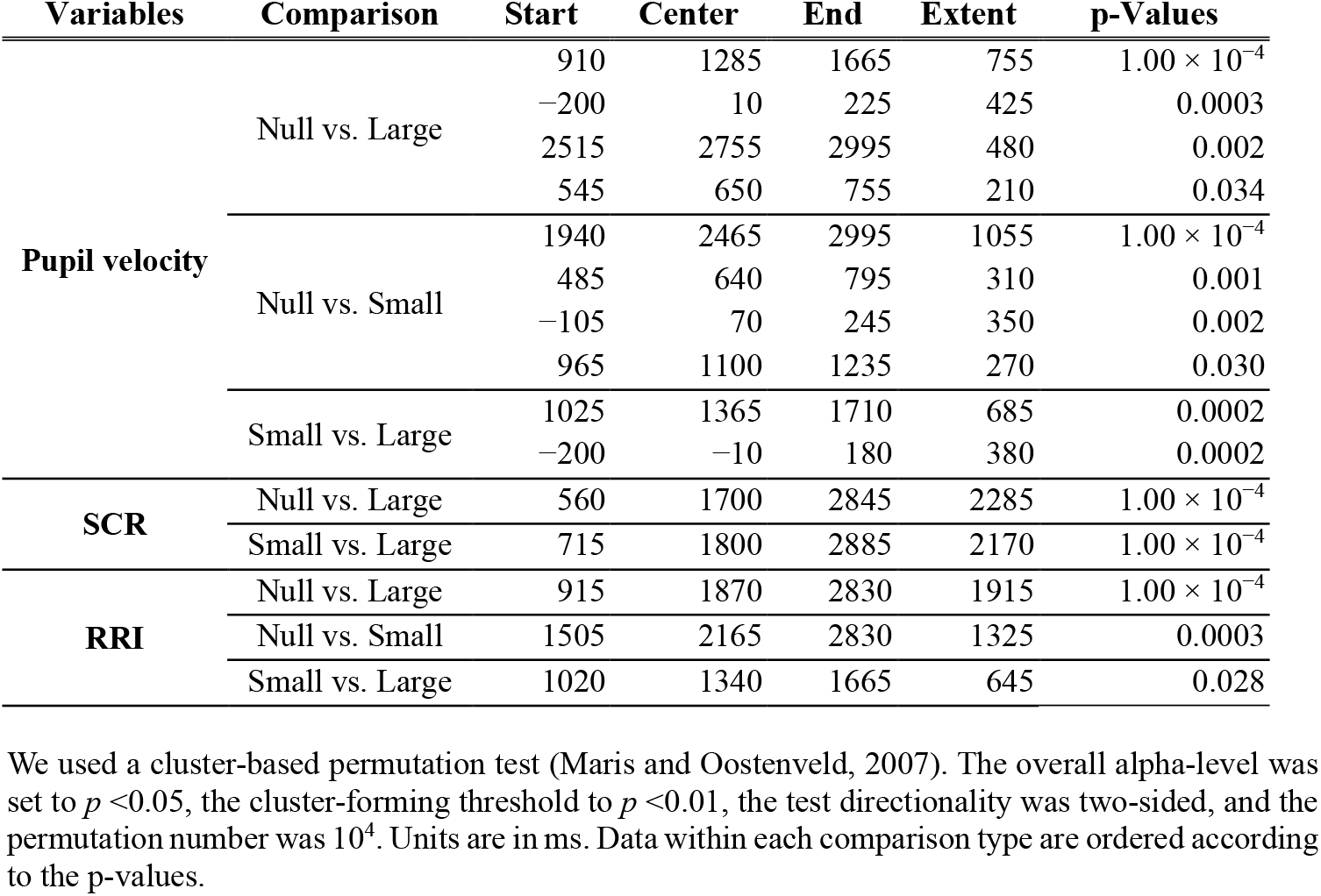
Temporal ROIs for the significant effect of proprioceptive perturbation (Exp 1) tested over the phasic autonomic response time series aligned to movement off (Fig. 2, right column).

**Table 3.**
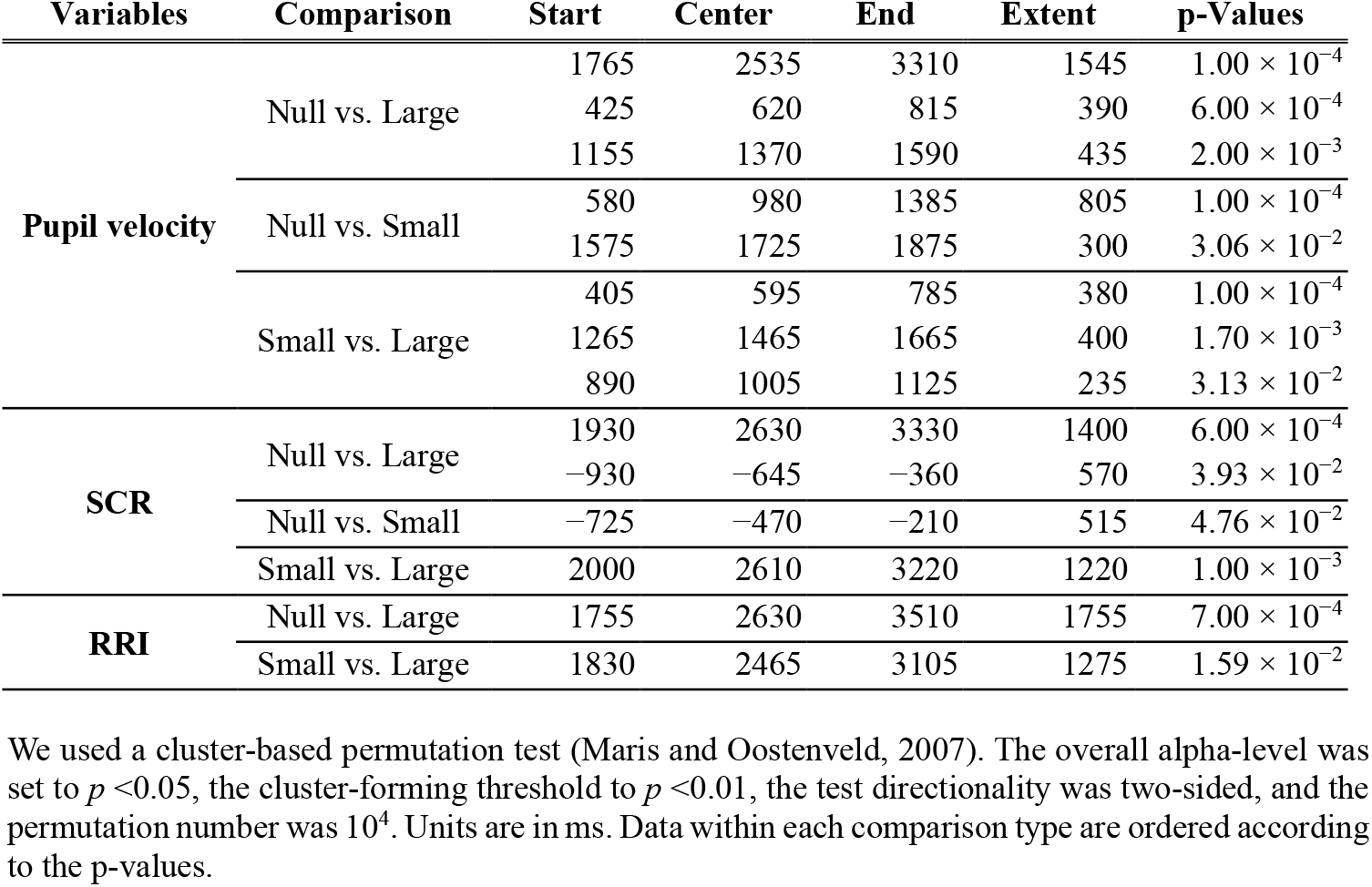
Temporal ROIs for the significant effect of visual perturbation (Exp 2) tested over the phasic autonomic response time series aligned to movement onset (Fig. 4, left column).

**Table 4.** Temporal ROIs for the significant effect of the visual perturbation (Exp 2) tested over the phasic autonomic response time series aligned to the movement offset (Fig. 4, right column).

| Variables | Comparison | Start | Center | End | Extent | p-Values |
| --- | --- | --- | --- | --- | --- | --- |
| **Pupil velocity** | Null vs. Large | 960 | 1730 | 2500 | 1540 | $1.00 \times 10^{-4}$ |
| | | 485 | 605 | 730 | 245 | $4.57 \times 10^{-2}$ |
| | Null vs. Small | 135 | 465 | 795 | 660 | $1.00 \times 10^{-4}$ |
| | | 875 | 1110 | 1345 | 470 | $1.00 \times 10^{-4}$ |
| | Small vs. Large | 1815 | 1985 | 2160 | 345 | $9.00 \times 10^{-3}$ |
| | | 325 | 455 | 590 | 265 | $2.94 \times 10^{-2}$ |
| **SCR** | Null vs. Large | 995 | 1690 | 2390 | 1395 | $1.00 \times 10^{-4}$ |
| | Small vs. Large | 1190 | 1770 | 2350 | 1160 | $9.00 \times 10^{-4}$ |
| **RRI** | Null vs. Large | 860 | 1845 | 2830 | 1970 | $5.00 \times 10^{-4}$ |
| | Small vs. Large | 1000 | 1435 | 1870 | 870 | $1.55 \times 10^{-2}$ |

We used a cluster-based permutation test (Maris and Oostenveld, 2007). The overall alpha-level was set to $p < 0.05$ , the cluster-forming threshold to $p < 0.01$ , the test directionality was two-sided, and the permutation number was $10^4$ . Units are in ms. Data within each comparison type are ordered according to the p-values.

The summary values of ANS responses to errors were derived by calculating the difference in the trial-by-trial ANS response summary for each temporal ROI between perturbation and pre-perturbation trials. The resultant delta-pupil dilation velocities, delta-SCR, and delta-zRRI data were then used for statistical analysis.

### Statistical analyses

*Timeseries comparison*. To detect significant effects of the perturbation size (small vs. large) in the autonomic response timeseries data, we applied the cluster-based permutation test (Maris and Oostenveld, 2007). The overall alpha-level was set to *p* <0.05, the cluster-forming threshold to *p* <0.01, the test directionality was two-sided, and the permutation number was 10^4^. We used the Matlab implementation of the test by E. Gerber (permutest.m; https://www.mathworks.com/matlabcentral/fileexchange/71737-permutest). In addition, we employed a mass *t*-test, a simple series of *t*-tests on each time frame corrected for the false discovery rate (*q* < 0.05) (Benjamini and Hochberg, 1995). For this, we used the Matlab implementation of the correction by D. Groppe (fdr_bh.m; https://www.mathworks.com/matlabcentral/fileexchange/27418-fdr_bh). For our experiments, the two tests yielded similar results (Yokoi and Weiler, 2022).

### Structural equation modeling (SEM) modeling

To assess the relationship between the phasic autonomic responses to errors and the single-trial learning rate, we applied SEM. We defined a path diagram based on the hypothesis that autonomic responses affect the learning rate (Fig. 6E). As manifest (observed) variables, we used the autonomic response summary data (delta-pupil dilation velocities, delta-SCR, and delta-zRRI), the error data, and the single-trial learning rate data, averaged within each perturbation size. As latent (construct) variables, we defined the sympathetic activity state (“ *Sym*”), the parasympathetic activity state (“ *Para*”), the prediction error (or could be called surprise) (“ *PE*”), and the learning rate (“ *LR*”) (Fig. 6E). “ *PE*’ was assumed to be formative and the other latent variables to be reflective. The path diagram was defined based on physiological evidence and SEM modeling requirements. We assumed that only “ *Para*” affects the RRI index as the kinetics of the sympathetic drive on the heart rate is much slower than the parasympathetic drive (Berger et al., 1989; Ng et al., 2001). Similarly, we assumed that the SCR index was only directly affected by “ *Sym*,*”* as the sweat grands are solely innervated by the sympathetic terminals (Dawson et al., 2007). While pupil diameter is regulated by both sympathetic and parasympathetic pathways, we assumed that the pupil indices were directly affected by “ *Sym*” only. The effect from the parasympathetic pathway was expressed as a path from “ *Para*” to “ *Sym*.” Although this path also affects the SCR index, we used the current path diagram because more complex diagrams with separate SCR and pupil indices yielded path coefficients inconsistent with known physiological evidence that pupil dilation/constriction is regulated by sympathetic/parasympathetic activity (Supplementary Material). As our latent variables contained one formative construct (“ *PE*”) and data are not normally distributed, we chose to apply the partial least square SEM (PLS-SEM) approach (Vinzi et al., 2010). The opposite path from “ *Sym*” to “ *Para*,” which makes the path diagram cyclic, was omitted to avoid loops in PLS-SEM modeling (Vinzi et al., 2010). To this end, we used the PLS-SEM Toolbox developed by M. Aria (https://www.mathworks.com/matlabcentral/fileexchange/54147-pls-sem-toolbox). The 95% confidence intervals for the factor loading and path coefficients were estimated through bootstrapping. We conducted participant-wise resampling 5000 times.

## Acknowledgements

We thank S. Kitazawa for helpful discussion. We also thank Saeko Kato for assistance in data collection. This work was supported by JSPS KAKENHI (JP22H03501 and JP23K24758) and JST FOREST program (JPMJFR234Z) awarded to A.Y. M.N. is supported by a fellowship from JST SPRING program (JPMJSP2138).

**Table 5.**
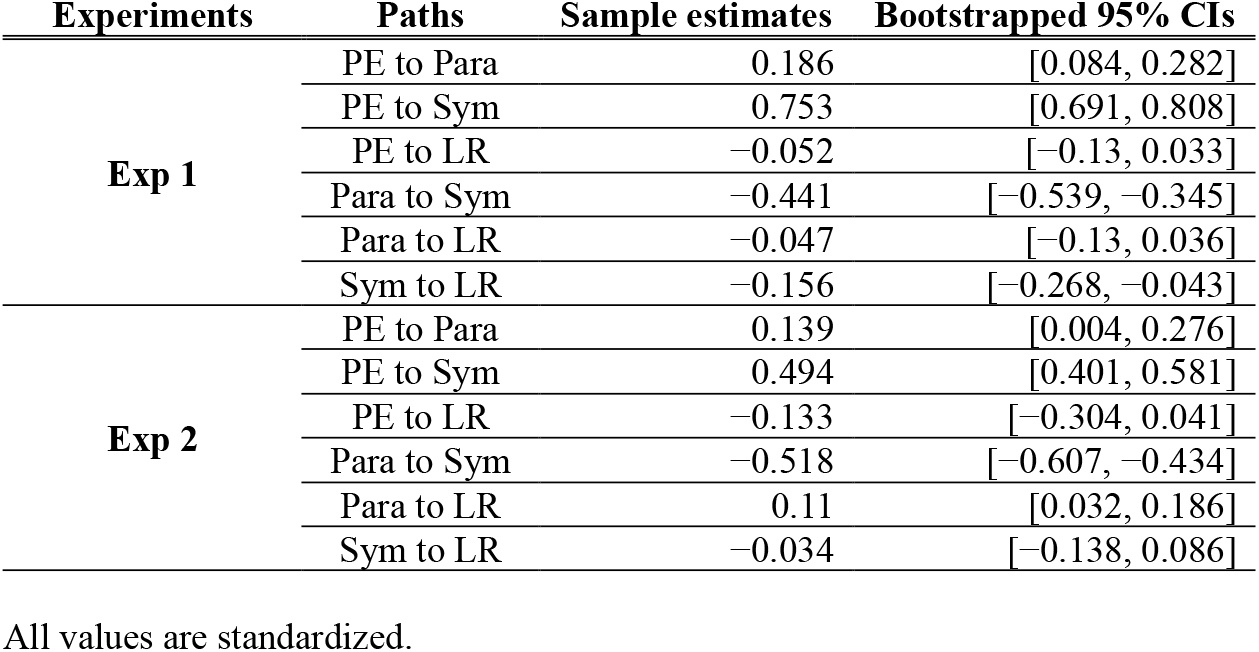
SEM results (path coefficients).

## Data and code availability

Data and custom-written Matlab codes used for analysis will be uploaded to a publicly available server upon publication.

## Results

To assess error-induced autonomic responses during motor learning, we simultaneously monitored the pupil diameter, SCL, and ECG while participants made reaching movements (Fig. 1A). In Exp 1 (n = 39), we manipulated the proprioceptive errors, while in Exp 2 (n = 39), we manipulated visual cursor errors, randomly selecting the error direction and magnitude among nine angles (±45°, ±30°, ±10°, ±5°, 0°; Fig. 1B, C). Given our interest in the ANS response to the motor error, as in previous research on the error-detection response in cognitive tasks (Hajcak et al., 2003; Preuschoff et al., 2011; Wessel et al., 2011; Nassar et al., 2012; Murphy et al., 2016), we only focused on the phasic error-induced response of ANS.

### Phasic autonomic responses to proprioceptive errors at distinct time windows (Exp 1)

We first evaluated the phasic change in the autonomic signal during the reaching task and their responses to movement errors. For pupil diameter and SCL data, we used the time derivative of the timeseries data to obtain the pupil dilation velocity and the SCR, to separate them from the tonic activity pattern. For ECG data, we evaluated the smoothed z-transformed RRI. Figure 2 shows the average time course of the pupil dilation velocity, SCR, and RRI in Exp 1, each aligned either to movement onset (left column) or offset (right column). As we did not provide any visual cursor or endpoint feedback during perturbation trials, the only error modality was proprioception.

**Figure 2.**
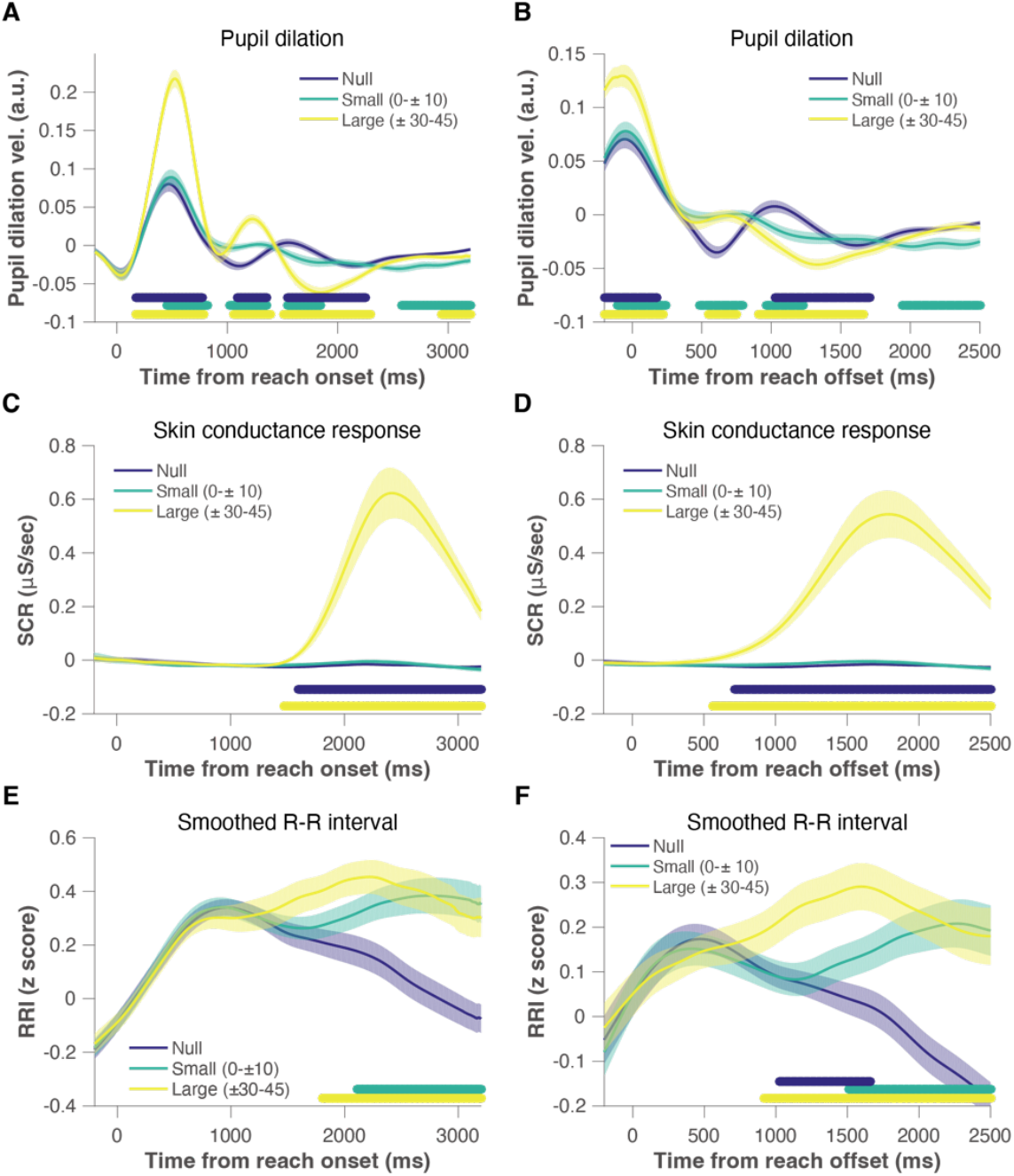
Time profiles of phasic autonomic responses to proprioceptive errors (Exp 1). **(A, C, E)** Phasic error responses measured in perturbation trials for pupil dilation velocity (A), skin-conductance response (C), and smoothed R-R interval (E) data aligned to movement onset time. **(B, D, F)** Phasic error responses for pupil dilation velocity (B), skin-conductance response (D), and R-R interval (F) data aligned to movement offset time. The shaded area represents the standard error of the mean (SE). Null: Normal reach trials interspersed between “ triplet” trials (Fig. 1B). Small: Perturbation trials with small proprioceptive errors (between −10° and 10°). Large: Perturbation trials with large proprioceptive errors (±30°, ±45°). The sequence of colored dots under the traces indicate significant (*p* < 0.05) clusters assessed by the cluster permutation test (Maris and Oostenveld, 2007). Blue: Comparison between Small vs. Large. Green: Small vs. Null. Yellow: Large vs. Null. Detailed statistics are presented in Tables 1 (onset-aligned data) and 2 (offset-aligned data).

During normal reaching trials (“ Null”), pupil (dilation) velocity and RRI showed the typical response observed in cue-response type tasks (van der Molen et al., 1989), whereas SCR showed a minor change (Fig. 2). When introducing the proprioceptive error by mechanically constraining the reach trajectory (Fig. 1B), these autonomic indices exhibited a clear phasic response to errors with specific latencies, and the response size was error size-dependent (Fig. 2). To extract the error-induced response components from the autonomic response time series, we first defined temporal ROIs by comparison between small (0°, ±5°, ±10°) (“ Small”) and large error trials (±30°, ±45°) (“ Large”). The average response data for the ROIs were later submitted to path analyses to test the potential effect of autonomic responses on LR.

The comparison between Small and Large conditions revealed three separate temporal ROIs for pupil dilation velocity with two dilation components followed by a constriction component (Fig. 2A; Table 1). The first error-induced dilation started as early as 170 ms after movement onset (Fig. 2A) and was due to proprioceptive error during movement. The other two ROIs were located after movement termination (Fig. 2A, B). For the SCR, the same comparison between “ Small” vs. “ Large” conditions revealed a clear increase in the SCR dependent on the error size at ∼1595 ms after movement onset (Fig. 2C). Considering the slow latency of the SCR (typically 1.0 ∼ 3.0 s from an event), this SCR is attributed to the proprioceptive error experienced during movement. Thus, it is likely that the initial error-induced pupil response was driven by sympathetic upregulation rather than parasympathetic withdrawal because sweat glands are predominantly innervated by sympathetic fibers (Dawson et al., 2007). Such sympathetic drive during movement was, however, not visible in the RRI, presumably due to the slower kinetics of sympathetic nerve effects on heart rate (Berger et al., 1989; Ng et al., 2001). Instead, the RRI responded to proprioceptive error in a parasympathetic direction (Fig. 2E, F). This RRI increase (i.e., heart rate deceleration) started after movement offset and showed a more pronounced effect when aligned to the movement offset (Fig. 2F, Table 2), suggesting a possible connection with postmovement evaluation/reflection. A previous study also reported the RRI increase after error commission in a choice reaction task (Danev and de Winter, 1971). Together, the autonomic measures we collected (pupil dilation velocity, SCR, and RRI) showed clear responses to unexpected proprioceptive errors and error size-dependency.

### Error size-dependent modulation of phasic autonomic responses to proprioceptive errors (Exp 1)

To look more closely into the error size-dependency of the autonomic responses, we analyzed the difference in the ANS responses between pre-perturbation and perturbation trials, for each error direction, size, and ROI (Fig. 3). As shown in Figure 3, the trial summary values of the phasic autonomic responses in each temporal ROI increased as a function of the error size, indexed by the amount of force received during the perturbation trials. The magnitude of proprioceptive errors was larger for rightward perturbations (defined as positive in Fig. 3), possibly because of the biomechanical properties (e.g., stiffness) of the right arm (Gomi and Kawato, 1996, 1997). Interestingly, visual inspection indicated that such asymmetry in the proprioceptive error for the same categorical perturbation sizes (e.g., +45° vs −45°) appeared to explain the size differences in the phasic autonomic responses (Fig. 3). Thus, the phasic autonomic responses seem sensitive to the discrepancy between the expected and actual force to the hand in a dose-dependent manner. In summary, the current results demonstrated that phasic pupil dilations followed by a constriction, SCR, and postmovement heart rate deceleration track the proprioceptive error size during movement.

**Figure 3.**
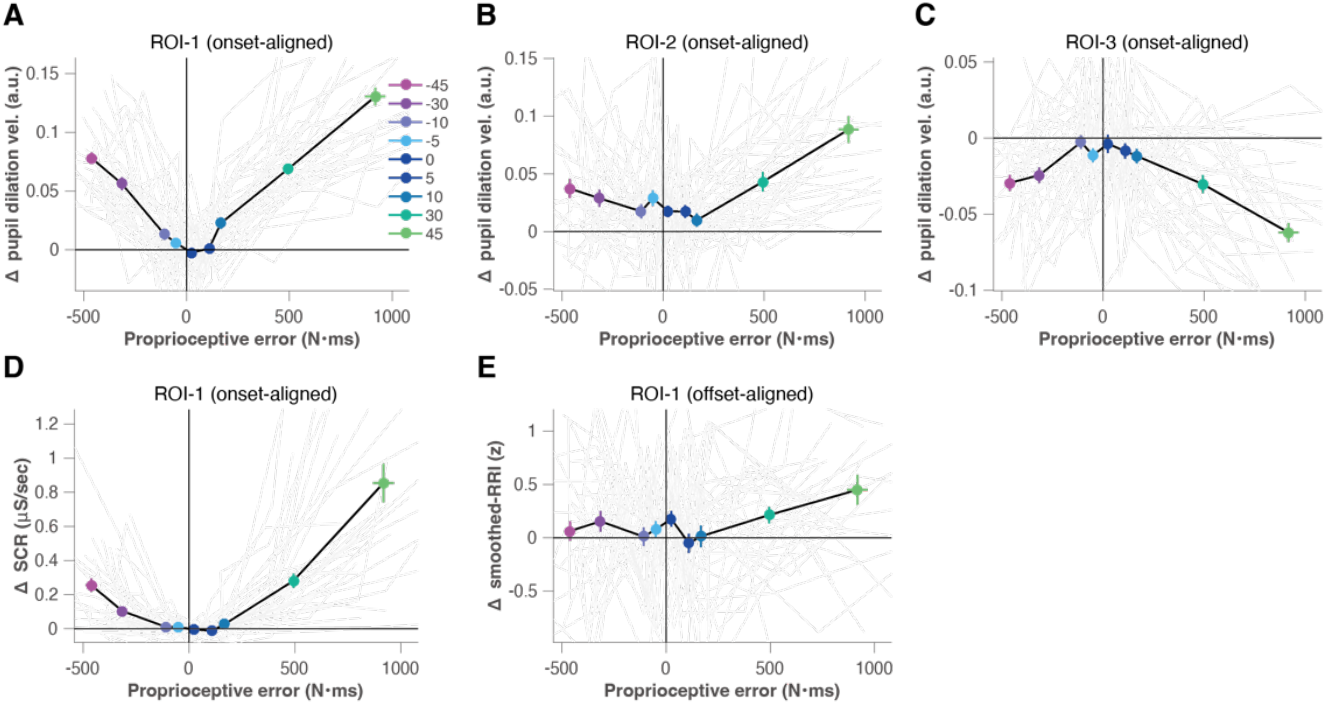
Mean phasic autonomic responses as a function of the proprioceptive error magnitude. Temporal regions of interest (ROIs) defined by the significant clusters shown in Fig. 2. The mean phasic autonomic signal data within each ROI were compared between perturbation vs. pre-perturbation trials to yield the delta-response data. **(A–C)** Mean delta-pupil dilation velocity for ROI-1 (A), ROI-2 (B), and ROI-3 (C), respectively (Fig. 2A). **(D)** Mean delta-SCR for ROI-1 (Fig. 2C). (E) Mean delta-smoothed-RRI for ROI-1 (Fig. 2F). Error-bars indicate the SEs for each pair of x-y values within each perturbation condition. Thin gray lines represent individual data.

### Phasic autonomic responses to visual errors (Exp 2)

To investigate the consistency of the ANS response to other error modalities, in Exp 2 we applied visual errors by rotating the cursor trajectory (i.e., visuomotor rotation) in a new set of participants (Fig. 1C). In contrast to Exp 1, we provided continuous visual cursor feedback, except for the post-perturbation trials where learning was probed in the no-feedback condition. This is to make the two experiments comparable by making online error information during reaching movements available. We also did not provide the endpoint feedback during the perturbation trial for the same comparability purpose.

In absence of perturbation (“ Null”), pupil (dilation) velocity, SCR, and RRI showed a similar response time course to that in Exp 1 (Fig. 2 and 4). For the error response, despite the weak magnitude, we found a qualitatively similar pattern of pupil and SCR responses, ensuring consistency across different error modalities (Fig. 4A–D). However, the first temporal ROI for the significant error size effect (“ Small” vs. “ Large”) in pupil dilation shifted in time (170 ms vs. 405 ms; Fig. 4A; Table 3). When we compared the range of the phasic response for the pupil and SCR in the first ROI across experiments, the response range for Exp 2 roughly matched that of Exp 1 for “ Small” perturbation (Fig. 3 and 5). Otherwise, the pupil response range for other ROIs and RRI change was comparable across experiments (Fig. 4). Overall, despite the weak magnitude of the error response in some measures in Exp 2, the autonomic measures showed qualitatively similar responses in terms of error size-dependency to Exp 1.

**Figure 4.**
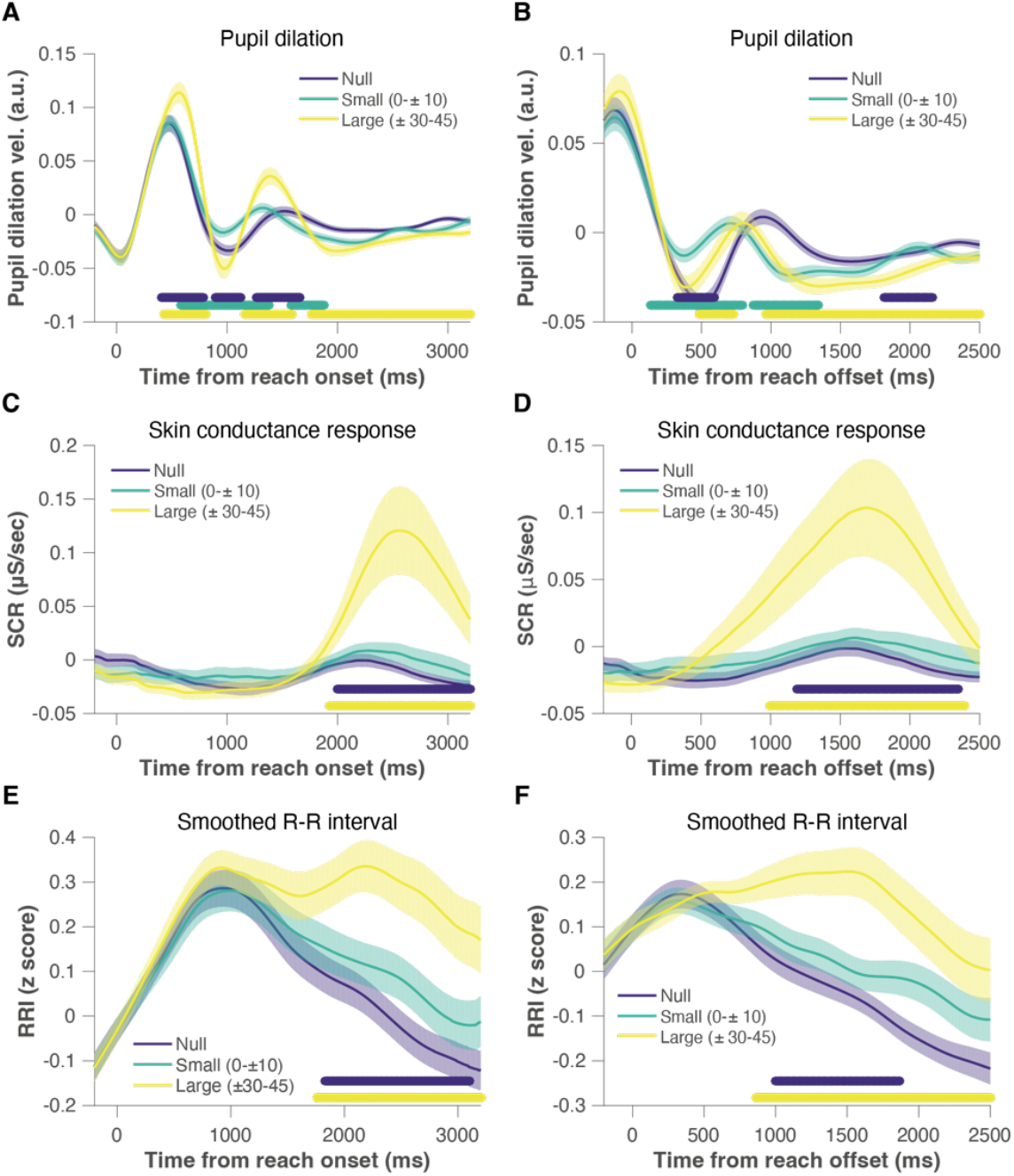
Time profiles of phasic autonomic responses to visual errors (Exp 2). **(A, C, E)** Phasic error responses measured at perturbation trials for pupil dilation velocity (A), skin-conductance response (C), and smoothed R-R interval (E) data aligned to movement onset time. **(B, D, F)** Phasic error responses for pupil dilation velocity (B), skin-conductance response (D), and R-R interval (F) data aligned to movement offset time. The shaded area represents the SE. Null: Normal reach trials interspersed between “ triplet” trials (Fig. 1B). Small: Perturbation trials with small proprioceptive errors (between −10° and 10°). Large: Perturbation trials with large proprioceptive errors (±30°, ±45°). The sequence of colored dots under the traces indicate significant (*p* < 0.05) clusters assessed by the cluster permutation test (Maris and Oostenveld, 2007). Blue: Comparison between Small vs. Large. Green: Small vs. Null. Yellow: Large vs. Null. Detailed statistics are presented in Tables 3 (onset-aligned data) and 4 (offset-aligned data).

**Figure 5.**
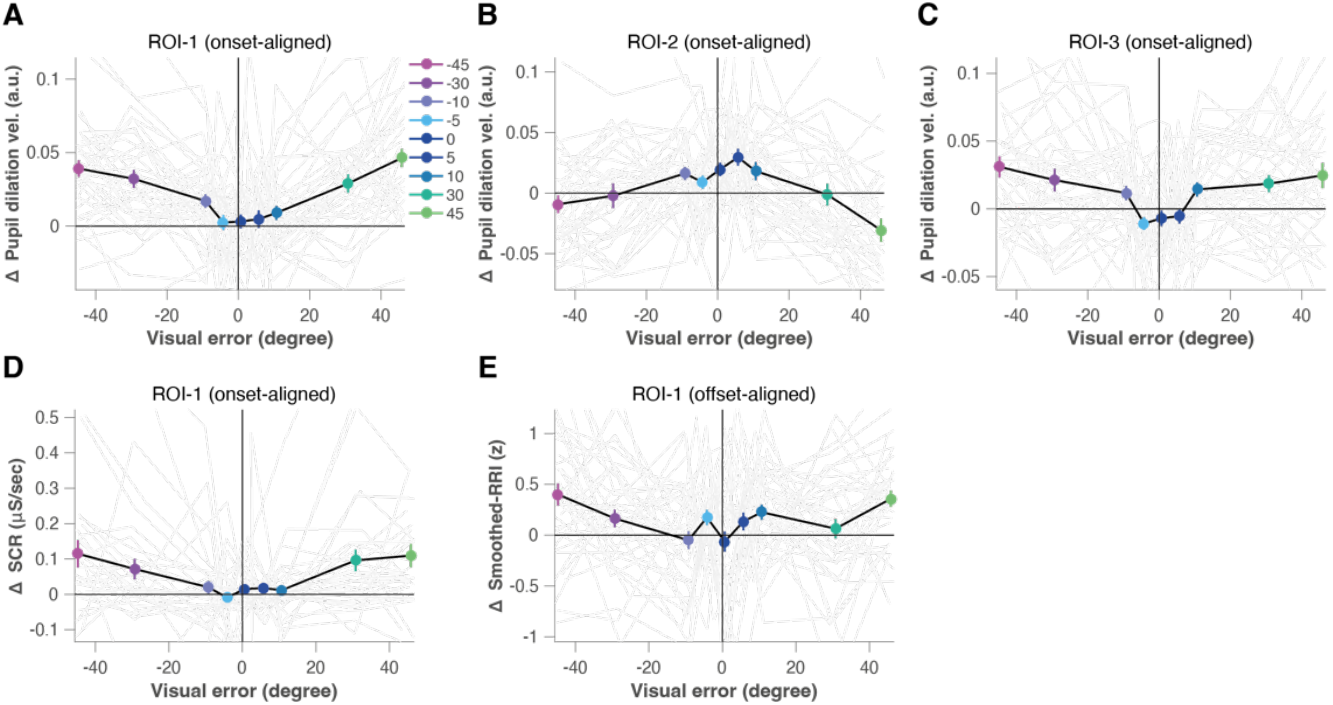
Mean phasic autonomic responses as a function of the visual error magnitude. Temporal regions of interest (ROIs) defined by the significant clusters shown in Fig. 4. The mean phasic autonomic signal data within each ROI were compared between perturbation vs. pre-perturbation trials to yield the delta-response data. **(A–C)** Mean delta-pupil dilation velocity for ROI-1 (A), ROI-2 (B), and ROI-3 (C), respectively (Fig. 4A). **(D)** Mean delta-SCR for ROI-1 (Fig. 4C). (E) Mean delta-smoothed-RRI for ROI-1 (Fig. 4F). Error-bars indicate the SEs for each pair of x-y values within each perturbation condition. Thin gray lines represent individual data.

**Figure 6.**
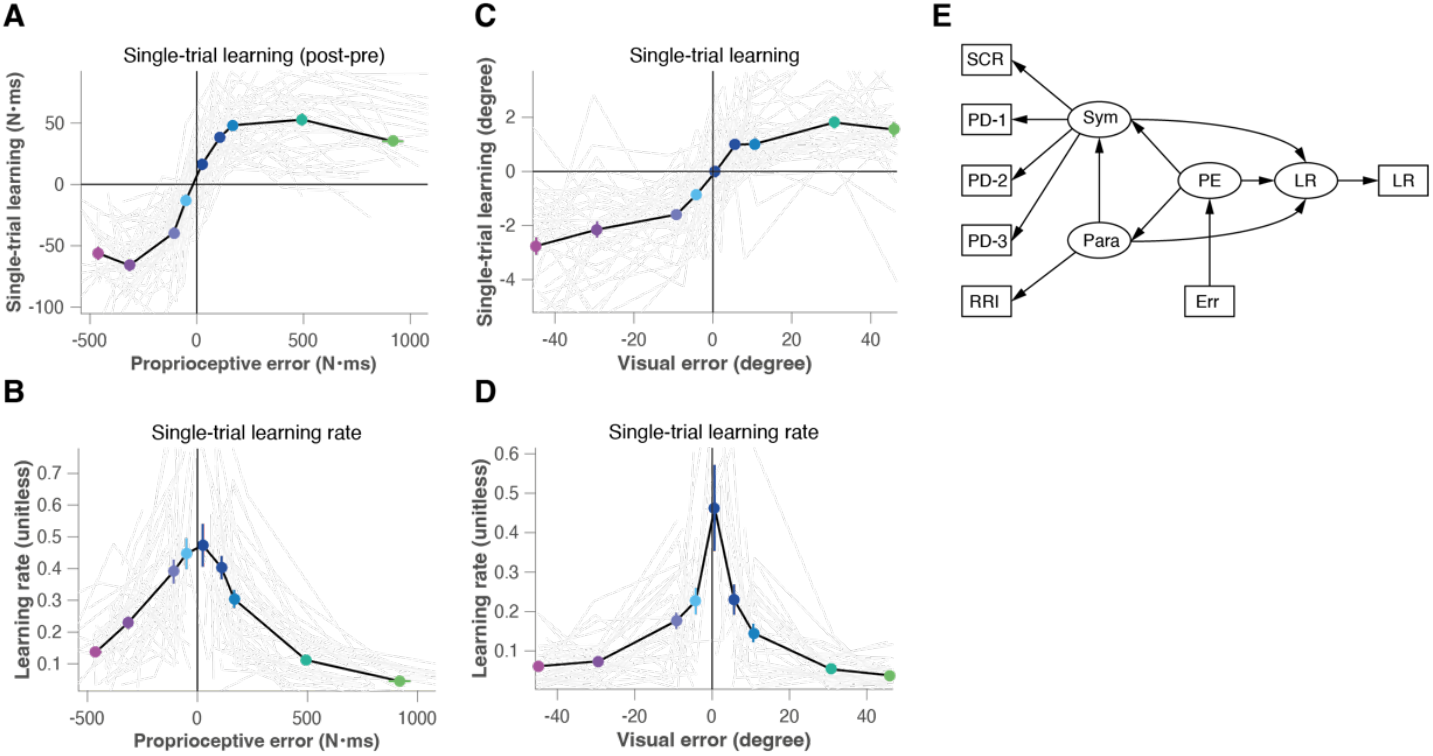
Single-trial motor learning and latent autonomic states. **(A, C)** Single-trial learning (across-trial correction) as a function of the proprioceptive (A) or visual (C) error size. The linear increase in correction only holds for small errors. **(B, D)** Single-trial learning rate as a function of the proprioceptive (B) or visual (D) error size. **(E)** Path diagram for the hypothetical relationship across observed (rectangles) and latent (ellipses) variables. SCR: Averaged SCR for each perturbation condition (Fig. 3D or Fig. 5D). PD-1–3: Averaged pupil dilation velocity for ROIs-1–3 for each perturbation condition (Fig. 3A–C or Fig. 5A–C). RRI: Averaged smoothed RRI for each perturbation condition (Fig. 3E or Fig. 5E). Sym: Latent state for sympathetic activity. Para: Latent state for parasympathetic activity. PE: Latent state for the surprise (unsigned prediction error). LR: Latent state for the learning rate. Error-bars indicate the SEs for each pair of x-y values within each perturbation condition. Thin gray lines represent individual data. The estimates and bootstrapped 95% CIs for the path coefficients and loadings are presented in Tables 5 (path coefficients) and 6 (loadings).

### Possible association between phasic autonomic error responses and the single-trial learning rate

Thus far, we observed that the magnitude of phasic autonomic responses increased with increasing induced movement error size. Is there any functional role for such modulation of the autonomic error response in motor learning? Interestingly, as reported in previous studies, the learning rate (sensitivity to errors) typically decreases for large errors compared to small errors (Fine and Thoroughman, 2006; Wei and Körding, 2009; Marko et al., 2012; Kasuga et al., 2013; Hayashi et al., 2020). Similarly, awareness to the perturbation depends on the perturbation size (Werner et al., 2015) which reportedly suppresses the implicit motor learning (Kagerer et al., 1997; Sakaguchi et al., 2001; Benson et al., 2011; Neville and Cressman, 2018). Consistently, such a pattern was present in our data (Fig. 6A–D). Across-trial corrections for larger errors were relatively smaller than for smaller errors (Fig. 6A, C), and sensitivity to error decreased with increasing error size (Fig. 6B, D). Based on these observations, we hypothesized that the error-induced phasic autonomic activity may be related to learning suppression depending on the error size.

To test this hypothesis, we evaluated the effect of phasic autonomic error responses on the rate of motor learning. Rather than comparing each autonomic signal to the learning rate separately or simply applying multiple linear regression, both susceptible to high measurement errors in predictor variables (i.e., autonomic indices in our case) and their correlation, we used SEM, which is robust to those problems by explicitly modeling the true underlying latent variables and their realization as noisy observations (Vinzi et al., 2010). Using SEM also allows us to combine multimodal ANS indices to obtain a more reliable estimate of an error-induced autonomic state and hence a more accurate estimate of the path between the autonomic state and the learning rate. We thus fitted a hypothetical relationship between the observed and latent variables (Fig. 6E). Here, we assumed three reflective latent states to express the fluctuation of the sympathetic (“ Sym”) and the parasympathetic (“ Para”) activities, as well as the learning rate (“ LR”) and one formative latent state for the internal representation of the unsigned prediction error (“ PE”), for each perturbation condition for each individual (Fig. 6E, see Methods for more detail). In this setting, the significant positive/negative path from the autonomic state (“ Sym” /” Para”) to “ LR” together with the significant path from “ PE” to “ Sym” /” Para” indicate the statistical mediation of learning rate modulation by the autonomic state.

The path model shown in Figure 6E fitted to Exp 1 data yielded a good fit; the standardized root mean square residual (SRMR) was <0.08 (= 0.0559) and the Tucker and Lewis index (TLI) >0.95 (=0.9668). As expected, the confirmatory factor analysis using SEM applied to Exp 1 data revealed a significant mediation effect by the error-induced sympathetic arousal on the suppression of motor learning rate. First, the path from “ Sym” to “ LR” was significant and negative (bootstrapped 95% confidence interval: [−0.268, −0.043]). Second, the path from “ PE” to “ Sym” was significant and positive (bootstrapped 95% confidence interval [0.691, 0.808]). Third, although the overall effect direction was negative, the path from “ PE” to “ LR” did not reach significance (bootstrapped 95% confidence interval: [−0.13, 0.033]). These results indicate the full-mediation effect of error-induced sympathetic activity in the error size-dependent suppression of the learning rate, supporting our hypothesis.

Next, we applied the same structural model to Exp 2 data to see if the result was consistent. This also yielded a decent fit (SRMR = 0.0539, TLI = 1.0196). However, despite the negative overall direction of the effect and the consistent results for “ PE” to “ Sym” and “ PE” “ LR” (see Tables 5 and 6 for complete results), the path from “ Sym” to “ LR” did not reach significance (bootstrapped 95% confidence interval: [−0.138, 0.086]). In summary, in spite of the different statistical level across experiments, our results point toward a consistent view, suggesting the possible involvement of error-induced sympathetic state changes in the modification of the trial-by-trial implicit motor learning rate^1^, in a direction of suppression.

**Table 6.**
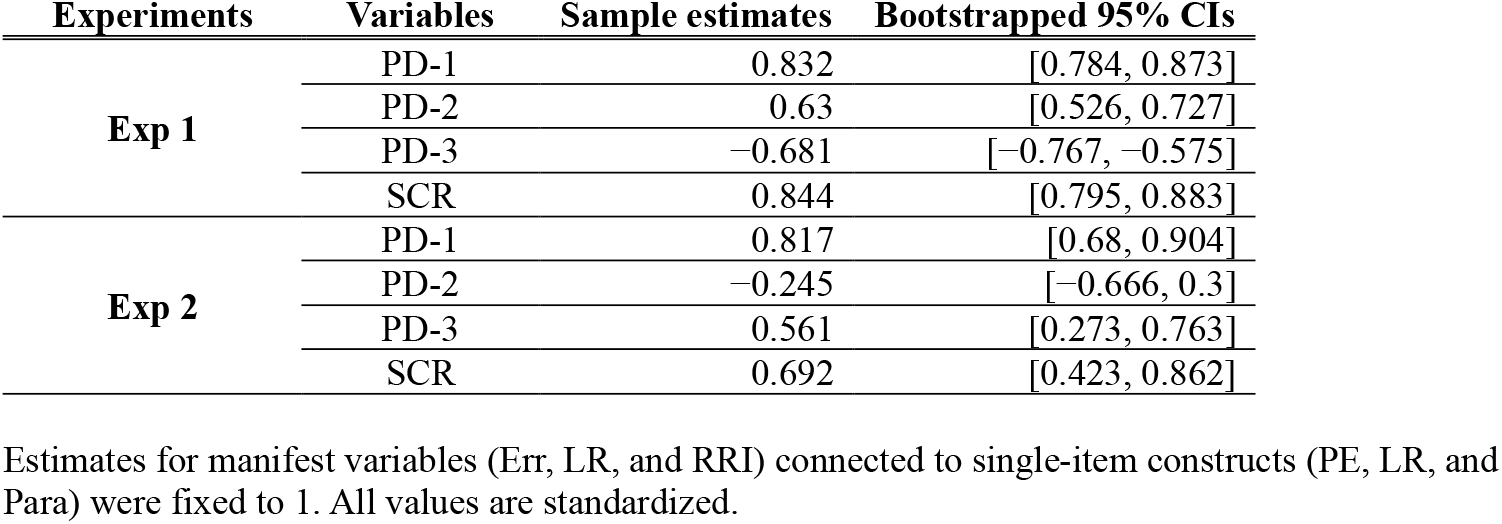
SEM results (loadings).

## Discussion

The current results captured the temporally dynamic nature of the multimodal ANS responses induced by unexpected errors during a motor learning task. These responses consisted of an initial sympathetic upregulation, as reflected by phasic pupil dilation and SCR, followed by parasympathetic dominance, as reflected by the heart rate deceleration. While the response magnitude, particularly for the initial sympathetic upregulation, was much greater in Exp 1, the ANS response patterns in the two experiments showed a qualitatively similar pattern; the response magnitude was scaled by the error size. In addition, the SEM analysis suggests that the autonomic states may mediate the error size-dependent adjustment of the implicit motor learning rate.

In our previous study (Yokoi and Weiler, 2022), we found phasic pupil dilation to unexpected reach errors caused by a sudden environmental change (i.e., force-field introduction). When the force field was gradually introduced to maintain the small size of the error, the error-induced pupil responses were minimal, suggesting an error size-dependent pupil dilation. Such tentative dose-dependency in the ANS error response was confirmed in the current study in a more direct and comprehensive manner through multimodal ANS measurement at different error sizes. Together with the recent view that phasic pupil dilation reflects a surprise-like internal state (Preuschoff et al., 2011; Nassar et al., 2012), our results suggest that ANS responses can be a continuous readout for such an internal state. This multimodal ANS measurement also confirmed that such a surprise-like state is indexed by sympathetic (LC-NA) activation rather than vagal withdrawal. Furthermore, the use of a single-trial learning paradigm allowed us to juxtapose the error-induced multimodal ANS responses and more accurate estimates of error sensitivity, compared to our previous report (Yokoi and Weiler, 2022). This allowed us to capture the negative association between the error-induced sympathetic state and implicit motor learning.

The sympathetic response in Exp 2, where the error was presented visually, was weaker than in Exp 1, where the error was presented proprioceptively. This result agrees with the fact that the visual input reaches LC more indirectly (Corneil and Munoz, 2014; Wang and Munoz, 2015; Joshi and Gold, 2020) compared to the proprioceptive input, which projects to LC via spinal projections (Szabadi, 2013; Poe et al., 2020). Moreover, during the error trial the visual cursor was presented in the peripheral vision with respect to the fixation point (the target was located 10 cm away from the starting position, which was approximately 13°), which may degrade the processing of error (Carrasco et al., 1995; Carrasco and Barbot, 2014; Zhang et al., 2024) and reduce the activity in the LC. Importantly, despite such potential differences (i.e., in the processing pathway and in the sensory uncertainty level), we consistently found a qualitatively similar ANS response pattern across experiments, i.e., the magnitude of phasic autonomic responses increased with increasing size of the motor error. This indicates the robustness of our findings.

We believe that the smaller magnitude of the ANS response in Exp 2 could explain why the influence (path) from the sympathetic system (Sym) to the learning rate (LR) did not reach significance, despite the matching sign of the influence (negative) between the two experiments. Nevertheless, future studies should aim to test this hypothesis.

The negative influence of the sympathetic (LC-NA) state on implicit motor learning observed in the current study fits well with known physiological evidence regarding the NA projection to the cerebellum (Bloom et al., 1971; Olson and Fuxe, 1971; Szabadi, 2013). Multiple studies have demonstrated that NA reduces the probability of complex spikes in cerebellar Purkinje cells, a key element for error-based motor learning (Kitazawa et al., 1998; Medina and Lisberger, 2008; Herzfeld et al., 2018), through the alpha2-adrenergic receptor (Carey and Regehr, 2009; Sun et al., 2019). Additionally, in the cerebellum, the alpha2-adrenergic receptor is dominant at high NA concentrations, while the beta-adrenergic receptor is dominant at low NA concentrations (Wakita et al., 2017), indicating suppression of complex spikes under high NA activity. These previous studies together with ours warrant further investigation regarding the functional role of NA in motor learning.

The pattern of ANS error responses observed during the reach adaptation task resembles previously reported error-related ANS responses during cognitive control tasks (Danev and de Winter, 1971; Hajcak et al., 2003; Preuschoff et al., 2011; Wessel et al., 2011; Nassar et al., 2012; Murphy et al., 2016; Di Gregorio et al., 2024). The similarity in the ANS response may imply a similarity in cognitive processes and brain areas (i.e., ACC and anterior insular cortex, AIC) involved during both motor and cognitive tasks. However, the suppression of implicit motor learning by error-induced sympathetic (LC-NA) arousal contrasts with the positive relationship between the error-induced ANS responses and the subsequent behavioral adjustment in cognitive tasks (Hajcak et al., 2003; Nassar et al., 2012; Ullsperger et al., 2014; Murphy et al., 2016). Such a discrepancy may suggest a reciprocal interaction between explicit cognitive and implicit processes during motor learning. One possible scenario is that the experience of a large error elicits the cognitive control process to kick-in, regardless of whether the task is more cognitive or motor. When the task involves precise motor control, the cognitive control process may particularly suppress implicit motor learning to avoid any unwanted adaptation of the current internal model, to let the process (or new internal model) handle the exceptional case that induced the large error.

Accordingly, awareness of errors suppresses implicit motor learning as measured by aftereffects (Kagerer et al., 1997; Sakaguchi et al., 2001; Benson et al., 2011; Neville and Cressman, 2018). Importantly, the activity of ACC and AIC, which have projections to the LC and are involved in the regulation of other autonomic controllers (Botvinick et al., 2001; Critchley et al., 2003, 2005; Ullsperger et al., 2014), is modulated by errors and their awareness (Klein et al., 2007; O’Connell et al., 2007; Wessel et al., 2011). These observations collectively support our idea that ANS responses could indicate suppression of implicit motor learning due to cognitive involvement. However, this hypothesis warrants further studies that simultaneously assess the error awareness, explicit learning component, and autonomic measures.

The cognitive intervention in the implicit motor learning system could be considered an instantiation of the recursive theory of contextual inference in motor learning (Haruno et al., 2001; Heald et al., 2021, 2023), where the “ upper” contextual inference process governs the formation and retrieval of “ subordinate” implicit motor memory. As repeatedly suggested in previous studies (Bouret and Sara, 2005; Sara, 2009; Sara and Bouret, 2012), the LC-NA system is in an ideal position to detect a contextual change through a large prediction error and transmit it to the memory systems throughout the brain. Thus, as a peripheral readout of the LC-NA system, ANS could provide an informative window for probing the latent computational variables relevant for such a contextual inference process.

In summary, the current study systematically characterized the changes in pupil diameter, SCR, and RRI in response to sudden errors during motor adaptation. Our results suggest that a closer look into the autonomic response during motor learning may help us better understand the interplay between cognitive and implicit learning processes underlying sophisticated motor skills.

## Author contributions

A.Y. conceived and designed the research; A.Y. and M.N. performed the experiments; A.Y. and M.N. analyzed the data; A.Y. and M.N. interpreted the results of the experiments; A.Y. and M.N. prepared the figures and tables; M.N. drafted the manuscript; A.Y., M.N. and N.H. edited and revised the manuscript; A.Y., M.N. and N.H. approved the final version of the manuscript.

## Footnotes

1 We presume that we quantified the implicit motor learning rather than a mixture of implicit learning and explicit strategies. We instructed the participants to always aim at the center of the target and confirmed that most participants did so in the post-experiment questionnaire (36/39 and 35/39 participants for Exp1 and 2, respectively).

