## Supplementary Material for "Autonomic responses to proprioceptive and visual errors during single-trial reach adaptation"

### SEM analysis: Alternative path models

We fitted additional path models that can capture the fact that pupil diameter is regulated by both of sympathetic and parasympathetic tones. Figure S1 A shows one such path diagram with an additional latent state for pupil variables (“SymPara”) receiving paths from “Sym” and “Para” latent states. We fitted three path models with different degrees of redundancy in paths to the data (Fig. S1A-C). As shown in Table S1, the overall quality of fit was slightly worse than the main model (Fig. 6E). Notably, while the positivity of the paths from “PE” to “Sym”, “Para”, and “SymPara” was consistent with the main result (Table 5), the path from “Para” to “SymPara” was positive for all three models presented in Figure S1 (Tables S2, 4, and 6). The path from “Sym” to “SymPara” was weak and sometimes not significant (Tables S2, 4, and 6). In all models, the path from “PE” to “LR” did not reach to significnce. Overall, these models yielded path estimates that are inconsistent with known physiological evidence of pupillary regulation.

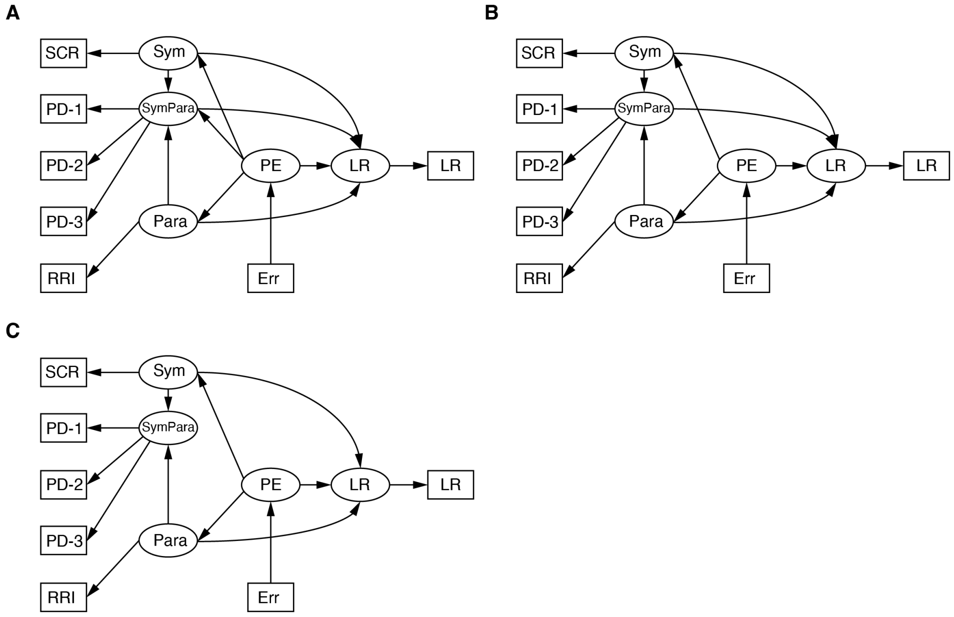


**Figure S1. Alternative path diagrams.**

**Table S1.** SEM results for alternative models shown in Figure S1 (fit summary).

| Experiments | Models | SRMR | TLI |
|---|---|---|---|
| **Exp 1** | A | 0.063 | 0.964 |
| | B | 0.065 | 0.957 |
| | C | 0.068 | 0.952 |
| **Exp 2** | A | 0.066 | 0.964 |
| | B | 0.075 | 0.948 |
| | C | 0.084 | 0.919 |

SRMR: standardized rootmean square residuals. TLI: Tucker and Lewis index.

**Table S2.** SEM results for the alternative model shown in Figure S1A (path coefficients).

| Experiments | Paths | Sample estimates | Bootstrapped 95% CIs |
|---|---|---|---|
| **Exp 1** | PE to Para | 0.186 | [0.084, 0.283] |
| | PE to Sym | 0.751 | [0.56, 0.751] |
| | PE to SymPara | 0.474 | [0.343, 0.611] |
| | PE to LR | –0.078 | [–0.155, 0.002] |
| | Para to SymPara | 0.343 | [0.182, 0.481] |
| | Para to LR | –0.456 | [–0.549, –0.361] |
| | Sym to SymPara | –0.055 | [–0.139, 0.034] |
| | Sym to LR | 0.061 | [–0.033, 0.154] |
| | SymPara to LR | –0.205 | [–0.324, –0.087] |
| **Exp 2** | PE to Para | 0.139 | [0.002, 0.273] |
| | PE to Sym | 0.252 | [0.15, 0.35] |
| | PE to SymPara | 0.35 | [0.103, 0.528] |
| | PE to LR | –0.021 | [–0.129, 0.094] |
| | Para to SymPara | 0.38 | [0.097, 0.623] |
| | Para to LR | –0.507 | [–0.592, –0.425] |
| | Sym to SymPara | 0.122 | [0.043, 0.198] |
| | Sym to LR | 0.095 | [–0.037, 0.248] |
| | SymPara to LR | –0.119 | [–0.262, 0.025] |

All values are standardized.

**Table S3.** SEM results for the alternative model shown in Figure S1A (loadings).

| Experiments | Variables | Sample estimates | Bootstrapped 95% CIs |
|---|---|---|---|
| **Exp 1** | PD-1 | 0.838 | [0.791, 0.877] |
| | PD-2 | 0.679 | [0.579, 0.769] |
| | PD-3 | –0.748 | [–0.822, –0.65] |
| **Exp 2** | PD-1 | 0.87 | [0.726, 0.959] |
| | PD-2 | –0.122 | [–0.675, 0.587] |
| | PD-3 | 0.639 | [0.35, 0.814] |

Estimates for manifest variables (Err, LR, SCR, and RRI) connected to single-item constructs (PE, LR, Sym, and Para) were fixed to 1. All values are standardized.

**Table S4.** SEM results for the alternative model shown in Figure S1B (path coefficients).

| Experiments | Paths | Sample estimates | Bootstrappeed 95% Cis |
|---|---|---|---|
| **Exp 1** | PE to Para | 0.186 | [0.083, 0.281] |
| | PE to Sym | 0.663 | [0.557, 0.75] |
| | PE to LR | –0.029 | [–0.125, 0.064] |
| | Para to SymPara | 0.648 | [0.576, 0.717] |
| | Para to LR | –0.458 | [–0.55, –0.367] |
| | Sym to SymPara | –0.055 | [–0.143, 0.032] |
| | Sym to LR | 0.06 | [–0.038, 0.156] |
| | SymPara to LR | –0.202 | [–0.317, –0.086] |
| **Exp 2** | PE to Para | 0.139 | [0.001, 0.272] |
| | PE to Sym | 0.252 | [0.147, 0.352] |
| | PE to LR | 0.024 | [–0.115, 0.171] |
| | Para to SymPara | 0.508 | [0.269, 0.681] |
| | Para to LR | –0.513 | [–0.591, –0.437] |
| | Sym to SymPara | 0.121 | [0.042, 0.198] |
| | Sym to LR | 0.105 | [–0.022, 0.263] |
| | SymPara to LR | –0.129 | [–0.266, 0.006] |

All values are standardized.

**Table S5.** SEM results for the alternative model shown in Figure S1B (loadings).

| Experiments | Variables | Sample estimates | Bootstrapped 95% CIs |
|---|---|---|---|
| **Exp 1** | PD-1 | 0.828 | [0.767, 0.875] |
| | PD-2 | 0.681 | [0.578, 0.772] |
| | PD-3 | –0.761 | [–0.833, –0.667] |
| **Exp 2** | PD-1 | 0.861 | [0.677, 0.975] |
| | PD-2 | 0.114 | [–0.487, 0.691] |
| | PD-3 | 0.662 | [0.36, 0.83] |

Estimates for manifest variables (Err, LR, SCR, and RRI) connected to single-item constructs (PE, LR, Sym, and Para) were fixed to 1. All values are standardized.

**Table S6.** SEM results for the alternative model shown in Figure S1C (path coefficients).

| Experiments | Paths | Sample estimates | Bootstrapped 95% CIs |
|---|---|---|---|
| **Exp 1** | PE to Para | 0.186 | [0.08, 0.284] |
| | PE to Sym | 0.663 | [0.558, 0.751] |
| | PE to LR | –0.035 | [–0.128, 0.063] |
| | Para to SymPara | 0.647 | [0.574, 0.715] |
| | Para to LR | –0.553 | [–0.635, –0.472] |
| | Sym to SymPara | –0.039 | [–0.124, 0.044] |
| | Sym to LR | –0.009 | [–0.11, 0.095] |
| **Exp 2** | PE to Para | 0.139 | [0.005, 0.267] |
| | PE to Sym | 0.252 | [0.145, 0.348] |
| | PE to LR | 0.013 | [–0.127, 0.162] |
| | Para to SymPara | 0.512 | [0.299, 0.677] |
| | Para to LR | –0.549 | [–0.614, –0.483] |
| | Sym to SymPara | 0.124 | [0.053, 0.199] |
| | Sym to LR | 0.05 | [–0.051, 0.165] |

All values are standardized.

**Table S7.** SEM results for the alternative model shown in Figure S1C (loadings).

| Experiments | Paths | Sample estimates | Bootstrapped 95% CIs |
|---|---|---|---|
| **Exp 1** | PD-1 | 0.815 | [0.734, 0.879] |
| | PD-2 | 0.701 | [0.59, 0.793] |
| | PD-3 | –0.762 | [–0.836, –0.664] |
| **Exp 2** | PD-1 | 0.815 | [0.531, 0.982] |
| | PD-2 | 0.239 | [–0.407, 0.784] |
| | PD-3 | 0.687 | [0.398, 0.913] |

Estimates for manifest variables (Err, LR, SCR, and RRI) connected to single-item constructs (PE, LR, Sym, and Para) were fixed to 1. All values are standardized.
